# Modelling the spatial crosstalk between two biochemical signals explains wood formation dynamics and tree-ring structure

**DOI:** 10.1101/2020.04.01.019638

**Authors:** Félix P. Hartmann, Cyrille B. K. Rathgeber, Eric Badel, Meriem Fournier, Bruno Moulia

## Abstract

In conifers, xylogenesis produces during a growing season a very characteristic tree-ring structure: large thin-walled earlywood cells followed by narrow thick-walled latewood cells. Although many factors influence the dynamics of differentiation and the final dimensions of xylem cells, the associated patterns of variation remain very stable from one year to the next. While radial growth is characterised by an S-shaped curve, the widths of xylem differentiation zones exhibit characteristic skewed bell-shaped curves. These elements suggest a strong internal control of xylogenesis. It has long been hypothesised that much of this regulation relies on a morphogenetic gradient of auxin. However, recent modelling works have shown that while this hypothesis could account for the dynamics of stem radial growth and the zonation of the developing xylem, it failed to reproduce the characteristic tree-ring structure. Here we investigated the hypothesis of a regulation by a crosstalk between auxin and a second biochemical signal, using dynamical modelling. We found that, in conifers, such a crosstalk is sufficient to simulate the characteristic features of wood formation dynamics, as well as the resulting tree-ring structure. In this model, auxin controls cell enlargement rates while another signal (e.g., cytokinin, TDIF) drives cell division and auxin polar transport.

**Highlight:** A dynamical model proves that two interacting signals (auxin, plus a cytokinin or the TDIF peptide) can drive wood formation dynamics and tree-ring structure development in conifers.

## Introduction

Tree radial growth relies on the production of new cells by the cambium and their subsequent differentiation. This process presents a high level of plasticity, contributing to the ability of trees to acclimate to changing environmental conditions (Ragni and Greb 2018). Therefore, in the current context of climate change, increasing attention is paid to the influence of the environmental factors on wood formation. However, the anatomical structure of conifer tree rings as revealed through tracheidograms, with their succession of large thin-walled earlywood cells and narrow thick-walled latewood cells, demonstrates a strikingly stable organisation under contrasting conditions (Balducci et al. 2016; Kiorapostolou et al. 2018; Cuny et al. 2018). Over one growing season, xylem radial growth generally follows a typical Gompertz curve, whose parameters depends on internal and external factors (Camarero et al. 1998; Rossi et al. 2003; Cuny et al. 2012). The monitoring of wood formation, through microcore samplings along the growing season, reveals that the developing xylem generally displays a zonation pattern composed of (1) a division zone (or cambial zone *sensu stricto)*, where cells grow and divide; (2) an enlargement zone, where cells grow without dividing; (3) a maturation zone, where non-growing cells undergo secondary wall deposition and wall lignification; and (4) a mature zone, composed of dead, fully functional xylem cells (Wilson 1984; Rathgeber et al. 2016). Over the growing season, the width of each zone follows a specific skewed bell-shape curve (Cuny et al. 2013, 2014, 2015; Balducci et al. 2016).

The stability of these dynamic patterns over the growing seasons, and of the resulting tree-ring structure, suggests a tight internal control of xylem development. This becomes manifest when bark strips are removed (Brown and Sax 1962; Li and Cui 1988) or when cambial cells are put into culture (Barnett 1978). Indeed, where spatial organisation disappears, growth becomes exponential, and a callus is generally formed. A polarity field is thus required to organise the developing xylem into radial cell file. It is generally considered that this field could be established through the flow of biochemical signals between the phloem and the xylem. Indeed, the role played by several signals in the control of wood formation is well-documented (see reviews in Fischer et al. 2019 and Buttò et al. 2020). The radial distribution of auxin, the most-studied phytohormone, has been measured in several species and at different times and positions inside the forming wood during the growing season (Tuominen et al. 1997; Uggla et al. 1996, 1998, 2001), revealing a concentration peak around the cambial zone that varies in amplitude during the season. Based on these observations, some authors put forward the “morphogenetic-gradient hypothesis”, according to which the graded concentration profile of auxin prescribes the width of each zone and, eventually, the final sizes of produced xylem cells (Sundberg et al. 2000; Bhalerao and Bennett 2003) by specifying the successive developmental identities of the cells (division and enlargement).

However, it has been shown through dynamical modelling that while the morphogenetic-gradient hypothesis accounts for the shape of the xylem growth curve and for the seasonal dynamics of the developing xylem zonation, it fails to explain the dimensions of the produced tracheids and the final structure of the annual ring (Hartmann et al. 2017). As long as it is assumed that a single signal sets both division and enlargement identities—the core of the morphogenetic-gradient hypothesis— tracheid dimension patterns do not follow the typical conifer tree-ring structure that is commonly observed. Another issue was the prediction of unrealistic regular spatial oscillations of high amplitudes in final cell sizes.

In parallel, several models of tree-ring formation have focused on carbon and water resources (Deleuze and Houllier 1998; Vaganov et al. 2006, 2011; Hölttä et al. 2010; Wilkinson et al. 2015; Drew and Downes 2015; Schiestl-Aalto et al. 2015). But they all aim to establish relationships between environmental conditions and radial growth, while paying little attention to the biological mechanisms involved at the cellular level. More recently, Cartenì et al. (2018) developed an original functional approach and proposed a mechanism linking the seasonal variations in sugar availability in the cambium to the anatomical structure of tree rings. This model convincingly reproduces the typical conifer tree-ring structure, but do not fully represent the biological mechanisms behind tracheid differentiation since primary and secondary wall deposition are not distinguished. Another strong limitation is that cells grow independently of each other, making the model unable to capture the coordination of the xylogenesis processes at the tissue scale.

While carbon and water availabilities are indispensable for wood formation, a growing body of experimental works points at the driving role played by hormones and peptides (Etchells et al. 2015; Immanen et al. 2016; Gursanscky et al. 2016; Brackmann et al. 2018; Han et al. 2018; Smetana et al. 2019). Moreover, the stability of wood formation patterns, despite fluctuating environmental conditions, suggests an intrinsic regulatory action through biochemical signals, that can be presumed to be less sensitive that photosynthesis or water transport. Other signals than auxin are involved, such as the small peptide TDIF from the CLAVATA family, which enters the cambium from the phloem and maintains vascular stem cells (Hirakawa et al. 2008; Etchells et al. 2015); or the plant hormone cytokinin, whose regulatory effect on cambial activity has been reported in aspen (Nieminen et al. 2008). But the full picture of this regulation remains unclear and dynamical models are needed to disentangle the role played by each signal. In the *Arabidopsis thaliana* root, for instance, Muraro et al. (2013, 2014) and el-Showk et. (2015) developed models of vascular patterning based on a finding by Bishopp et al. (2011) that a crosstalk between auxin and cytokinin specifies developmental zones. To reproduce maize leaf growth profiles, De Vos et al. (2020) integrated hormonal crosstalk into a model and predicted the existence of a signal produced in the mature part of the leaf. Such approaches will be instrumental in understanding xylogenesis, with the additional challenge that not only growth profiles and developmental zonation have to be explained, but also the cell size pattern typical of tree rings.

To investigate the potential of the crosstalk between two biochemical signals in controlling tree radial growth, wood formation, and tree-ring structure, we further developed the XyDyS modelling framework. XyDyS2 assigns xylem cell identity based on two interacting biochemical signals.

## Material and Methods

### Model description

#### Core of the XyDyS2 model

Taking advantage of the symmetry of the xylem tissue, we only consider a single radial file of differentiating cells (Fig. 1). The radial file is composed of cells that either differentiate into tracheids within a given growing season (possibly after one or several division cycles) or remain undifferentiated in the cambium at the end of the season. We focus on a single growing season and the formation of one tree ring. Spatially, the first boundary of the system (“the cambium boundary”) is the interface with the part of the cambium which differentiates into phloem. The second boundary (“the xylem boundary”) is the interface with the mature xylem produced during the previous year. Within a file, cells are indexed from *i = 1*, at the cambium boundary, to *i* = *N(t)*, at the xylem boundary, *N*(*t*) being the number of cells in the radial file at time *t*. Each cell is geometrically characterised by its radial dimension, called “length” *L_i_(t). L(t)* denotes the total length of the radial file at time *t*. For the initial condition, we suppose that there are initially *N*_0_ cambial cells in the radial file, all with the same length *L*_init_.

**Figure 1:**
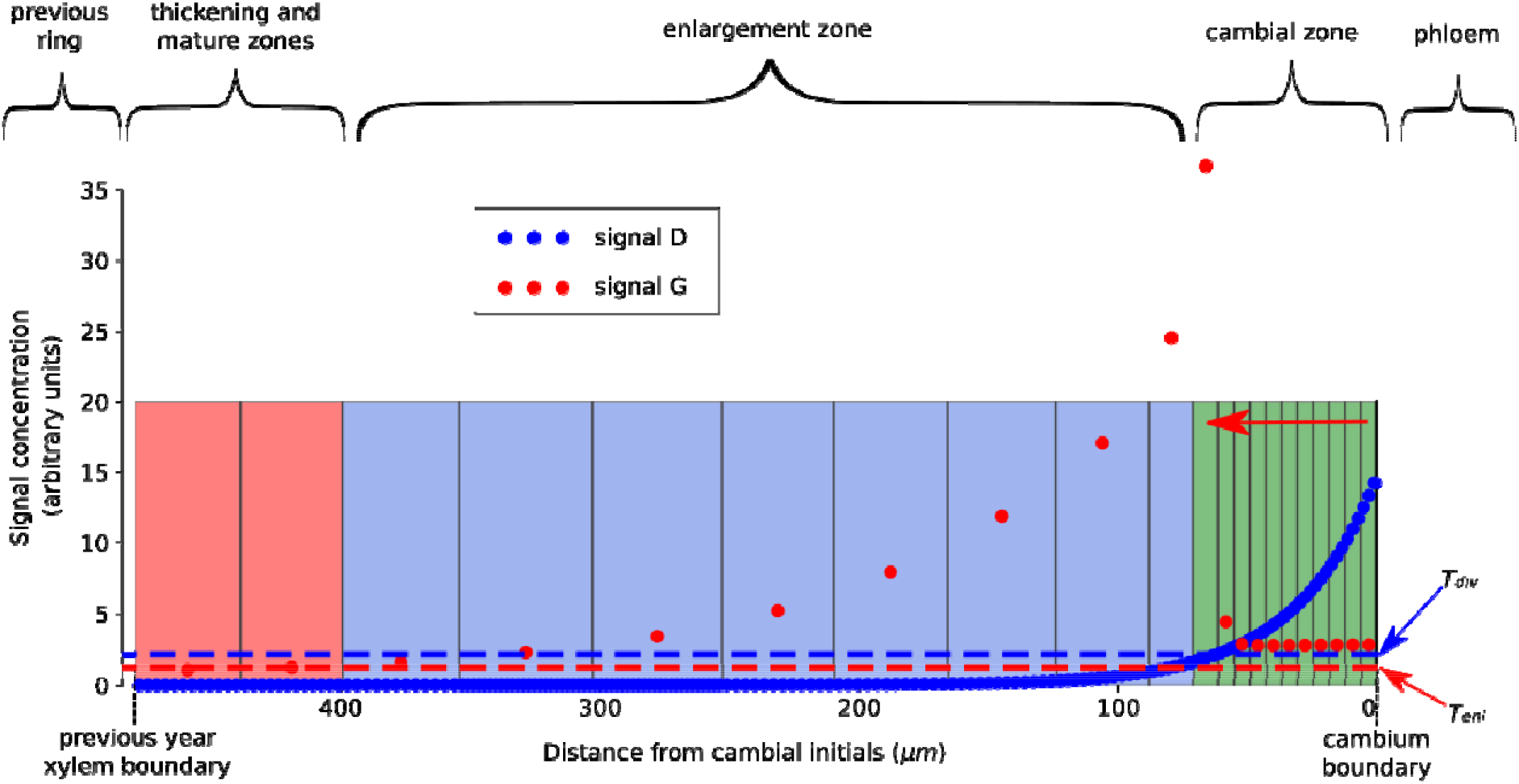
Schematic layout of a XyDyS simulation. Signals D and G form concentration gradients (respectively blue and red dots) which impose cell identities and growth rates. Cells with a concentration of signal D above the division threshold *(T_div_)* have the ability to divide. Cells with a concentration of signal G above the enlargement threshold (*T_enl_*) are growing, with a growth rate related to the concentration of signal G. Carrier proteins transporting signal G toward the xylem are present only in cells that have the ability to divide. The zonation is based on cell identity and geometry. Cambial zone (green): small (L_i_ < 2L_init_) growing cells. Enlargement zone (blue): large (L_i_ > 2L_init_) growing cells. Thickening zone and mature zone (red): non-growing cells.

Two signals, denoted by D and G, flow through the radial file, coming from the cambium boundary: Signal D is associated with cell division and could be identified as either the TDIF peptide or the cytokinin phytohormone, Signal G is associated with cell growth and is identified as auxin.

#### Apoplastic diffusion of signal D

The exact nature of the signal D is not elucidated, but we assume that it diffuses in the apoplast, like peptides and cytokinins do (Robert and Friml, 2009). The simplest model for signal diffusion is Fick’s law (Crick, 1970), with a constant decay rate. We also assume that signal D is not produced in the developing tissue but comes from an external source at the cambium boundary. This “source-diffusion-decay mechanism” is similar to that proposed by Wartlick et al. (2009) and Grieneisen et al. (2012) for root primary growth. Given the very slow growth of the developing xylem, dilution and advection (i.e. directed movement driven by tissue growth) can be neglected (Hartmann et al.

2017). Then, the transport equation of signal D writes as:

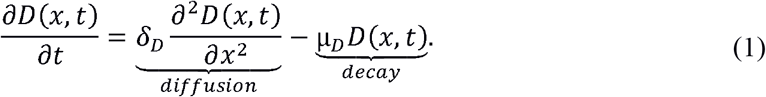

*D(x, t)* denotes the concentration of signal D at position *x* and time *t*, *δ_D_* denotes its diffusion coefficient and *μ_D_* its decay rate. The space variable *x* is defined such that the cambium boundary of the file is located at *x = 0* and the xylem boundary at *x* = *L(t).*

Equation 1 can be solved analytically. It is useful to introduce a characteristic length associated with the diffusion-decay process, expressed as:

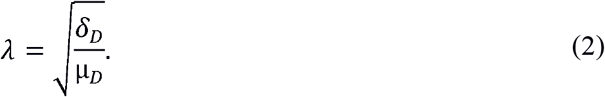

When the file becomes long compared to *λ*, the concentration profile reaches a stationary exponential shape, given by the equation:

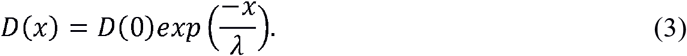

Finally, *D_i_* denotes the average concentration of signal *D* in cell *i*.

#### Symplastic polar transport of signal G

To describe the flow of signal G, identified as auxin (Perrot-Rechenmann 2010), we use a model of auxin fluxes similar to the “unidirectional transport mechanism” from Grieneisen et al. (2012) and Hartmann et al. (2017). Where PIN carrier proteins are present, auxin is polarly transported from one cell to another. In addition to this active transport, there is a residual constitutive permeability to auxin, which is the same between every consecutive cell. The auxin flux *F_i,i+1_* from cell *i* to cell *i+1* depends on the concentration of auxin in cell *i* and on the amount of PIN in cell *i* oriented toward cell *i+1* (Grieneisen et al. 2012). This writes as:

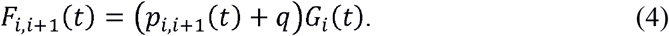

*G_i_(t)* is the concentration of signal G in cell *i, p_i,i+1_* is the amount of PIN proteins in cell *i* oriented toward cell *i+1*, and *q* is the constitutive permeability to auxin. Moreover, we assume that PIN proteins are always oriented toward the xylem, i.e. *p_i,i-1_ = 0*. Therefore, auxin fluxes toward the cambium boundary rely only on constitutive permeability, i.e. *F_i,i-1_(t)* = *qG_i_(t)*.

If one considers cell *i*, entering fluxes from cells *i-1* and *i+1* are respectively *F_i-1,i_* and *F_i+1,i_*, and exiting fluxes toward cells *i-1* and *i+1* are respectively *F_i,i-1_* and *F,_i,i+1_*. Considering also decay, and dilution due to cell growth, the concentration of signal G in cell *i* is governed by the following equation:

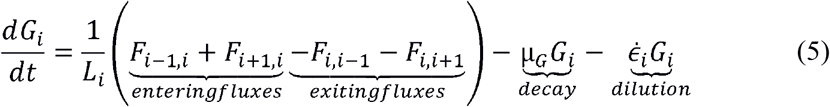

*μ_G_* is the decay rate of signal G, and 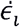 is the growth rate of cell *i* (Moulia and Fournier 2009), defined by:

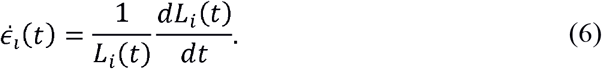

If fluxes are decomposed into polar and passive components, equation 5 becomes:

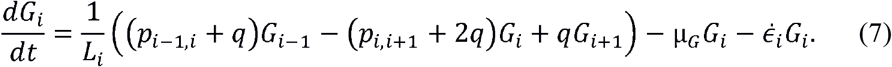

#### Cell identity assignment

In the classical morphogenetic-gradient model (Bhalerao and Fischer 2014; Hartmann et al. 2017), cell identities are set by a single signal, with two concentration threshold values: a division threshold *T_div_*, and an enlargement threshold *T_enl_*, with *T_div_ > T_enl_*. But this way of assigning identities leads to unrealistic patterns in mature tracheid diameters (Hartmann et al. 2017). Here, two distinct signals assign cell identities (Fig. 1). Where the concentration of signal D is higher than the division threshold *T_div_,*, cells are able to divide. Similarly, where the concentration of signal G is higher than the enlargement threshold *T_enl_*, cells enlarge. More formally, for a given cell:

- if *D_i_ ≥ T_div_*, the cell is able to enlarge and divide;
- if *D_i_ < T_div_* and *G_i_ ≥ T_enl_*, the cell is not able to divide anymore, but it can keep enlarging;
- if G*_i_* < *T_enl_*, the cell no longer enlarges.

Moreover, we assume that auxin efflux carriers (PIN proteins) are present only in cells that are able to divide (i.e. *p_i,i+1_ > 0* only if D*_i_*; ≥ *T_div_*). In these cells, the amount of PIN proteins, *p_i,i+1_*, is assumed to be proportional to the auxin concentration (signal G) in cell *i*:

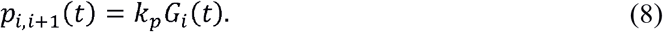

#### Cell growth and division

Although the mechanical force for cell enlargement comes from turgor pressure, this process is controlled by cell wall extensibility (Cosgrove, 2005). We assume that auxin acts on wall extensibility (Arsuffi and Braybrook 2018), and thus controls the growth rate of those cells which have an identity that allows them to enlarge. The simplest relationship is a direct proportionality: 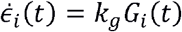, where k_g_ is a proportionality constant (Hartmann et al. 2017). However, this relationship implies exponential growth for constant levels of auxin, which tends to amplify inhomogeneities in cell sizes. Therefore, we propose here that larger cells display a weaker growth response to auxin, in the form of an inverse proportionality to cell size:

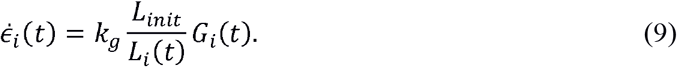

Cell division follows a simple geometrical criterion: if a cell has an identity that allows division, it divides when reaching a critical length defined as twice its initial length *L_init_* (Hartmann et al. 2017). All parameters of the model are listed in Table 1.

**Table 1:** Parameters of the model, with their value.

| Symbol | Value | Unit | Description |
| --- | --- | --- | --- |
| $N_0$ | 6 | unitless | Initial number of cells in the file. |
| $L_0$ | 6 | $\mu\text{m}$ | Initial size of the cells. |
| $T_d$ | 2 | unitless | Division threshold. |
| $T_e$ | 1.6 | unitless | Enlargement threshold. |
| $k_g$ | 0.06 | $\text{s}^{-1}$ | Prefactor relating signal concentration to cell growth rate. |
| $\delta_D$ | 10 | $\mu\text{m}^2 \cdot \text{s}^{-1}$ | Diffusion coefficient of signal D. |
| $\mu_D$ | $10^{-2}$ | $\text{s}^{-1}$ | Decay rate of signal D. |
| $\mu_G$ | $10^{-5}$ | $\text{s}^{-1}$ | Decay rate of signal G. |
| $q$ | $1.5 \times 10^{-3}$ | $\mu\text{m} \cdot \text{s}^{-1}$ | Permeability rate of the membranes to signal G. |
| $k_p$ | $4 \cdot 10^{-23}$ | unitless | Proportionality coefficient between the concentration of signal G in a cell and the amount of PIN in this cell. |

#### Definition of developing zones

Experimentally, the descriptions of the developmental zones are based on visual criteria. In order to be able to compare the outputs of the XyDyS2 model with real data, we apply similar criteria *a posteriori* on model outputs, setting “apparent statuses” to virtual cells:

- Cambial cells are growing cells that are smaller than two times the diameter of a newly created cell (*L_i_* < *2L_init_*).
- Enlarging cells are growing cells that are larger than two times the diameter of a newly created cell (*L_i_* > *2L_init_*).
- Wall-thickening and mature cells are no longer growing cells (Fig. 1).

#### Boundary conditions

The concentrations of signals D and G are imposed at the cambium boundary of the file, and are given as entries of the simulations (Fig. 2). The concentration of signal D at the cambium boundary is assumed to increase rapidly at the beginning of the season, and then decreases slowly. The cambium-boundary concentration of signal G is assumed to peak during the first weeks of the season, then progressively decrease to zero as the season goes. This reflects the sudden flush of auxin coming from the shoots during bud break. Finally, we assume that the xylem acts as an impermeable barrier to molecules of signals D and G. Accordingly, a zero-flux boundary condition is imposed for both signals at the xylem boundary.

**Figure 2:**
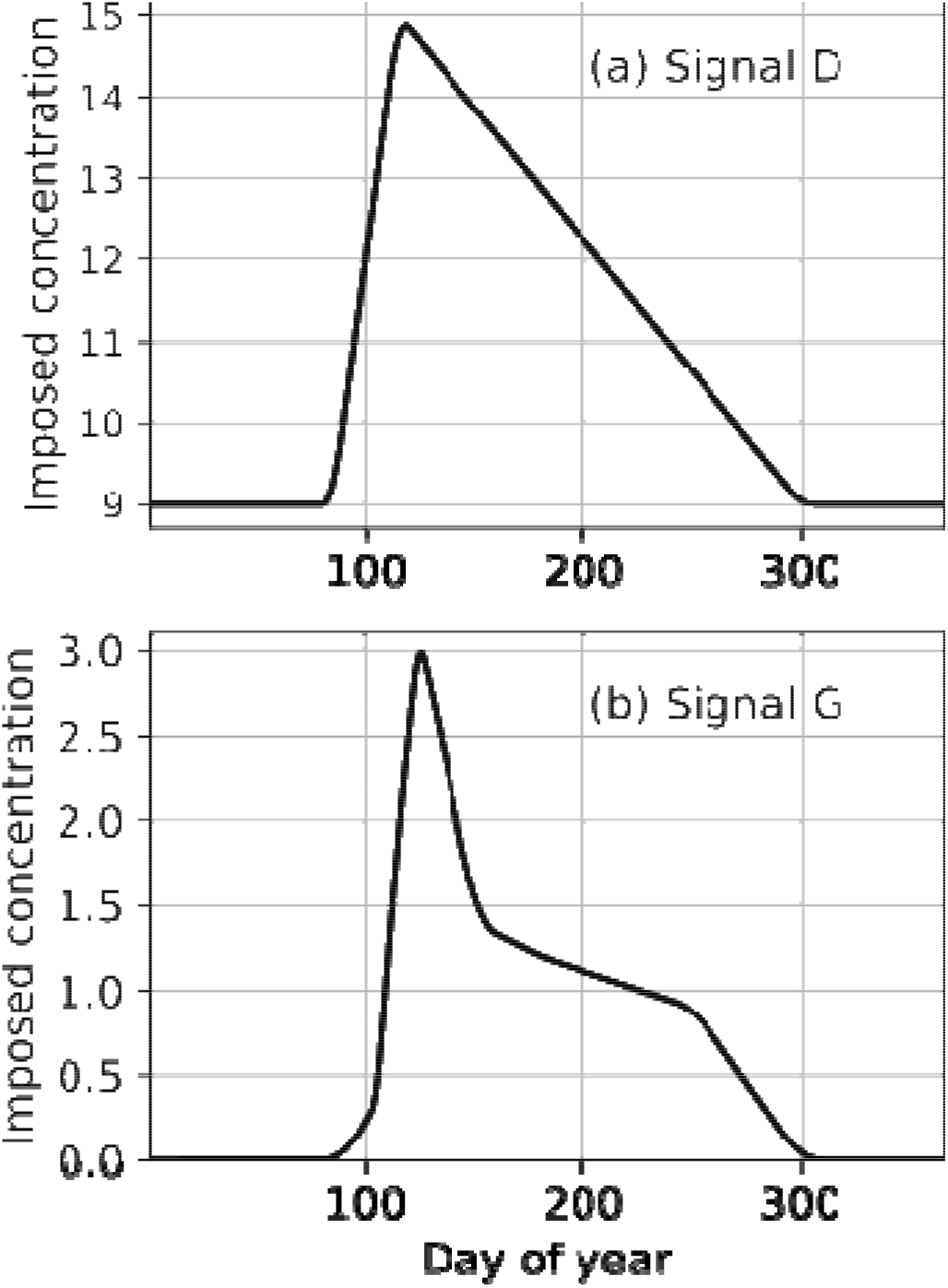
Concentrations of signals D and G imposed at the cambium boundary over a growing season.

#### Implementation and visualization of the simulations

Transport equations are numerically solved using an explicit Euler method. For signal D, which diffuses in the apoplast, additional discretisation nodes are regularly inserted in growing cells so that the Courant–Friedrichs–Lewy stability condition is always satisfied. We have developed a dedicated graphical user interface. The source code, written in Python, is freely available online (https://forgemia.inra.fr/felix.hartmann/xydys). Simulation outputs are visualized using the graphical convention explained in Fig. 1.

## Results

### The cross-talk between the two signals leads to the progressive establishment of a stable auxin gradient

We first looked at the establishment of signal concentration profiles at the beginning of the growing season. Since the length of the cell file was initially shorter than the characteristic length *λ*, signal D was filling in the cell file, with a high concentration everywhere (Fig. 3a and S1 Video). Therefore, all cells were dividing and transported auxin toward the xylem. As a consequence, signal G initially accumulated in the cells close to the xylem boundary, which thus had high growth rates. As the cell file became larger than *λ*, the concentration profile of signal D progressively reached the stationary exponential shape given by Eq. 3. From this time on, polar transport was limited to a few dividing cells and the concentration of signal G peaked around the boundary between the cambial and enlarging zones (Fig. 3b). The gradient of signal G was then stable and the height of the peak depended only on signal G concentration at the cambium boundary. Near the end of the growing season, the signal G became too low for a peak to form (Fig. 3c). This shows that active polar transport, regulated by another signal, can account for the peaked distribution of auxin observed experimentally in the developing xylem (Uggla et al. 2001).

**Figure 3:**
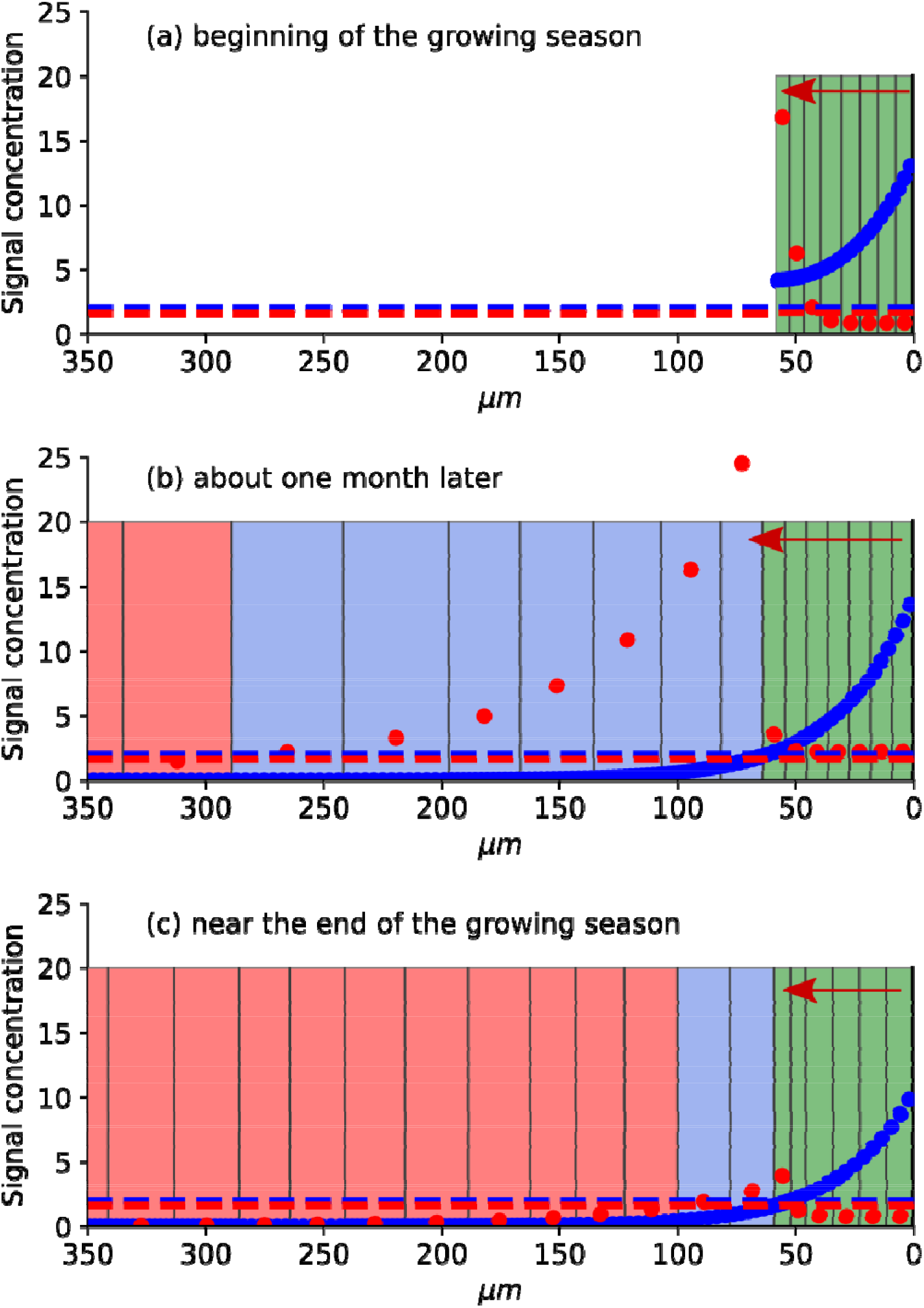
Establishment of signal gradients. (a) At the beginning of the growing season, signal D (blue dots) is above the division threshold in every cell. Signal G (red dots) is transported toward the xylem and accumulates at the xylem end of the cell file. (b) After the file has grown longer, both signals reach a stationary gradient. The concentration of signal G peaks around the boundary between the cambial and enlargement zones. (c) Near the end of the growing season, the supply of signal is very low.

### The cross-talk between the two signals controls the developmental zonation over the growing season

The width of the cambial zone was controlled mostly by signal D. At the beginning of the growing season, all cells belong to the cambium and the cambial zone expanded rapidly since signal D was high in every cell. This caused an early ‘burst’ in the number of cambial cells and, after a lag, in the number of enlarging cells. Such a rapid increase had also been observed in real wood formation monitoring studies (Cuny et al. 2014, 2018; Balducci et al. 2016). As the concentration profile of signal D stabilised into a stationary gradient, the number of cambial cells reached a constant value. This value depended only on the concentration of signal D imposed at the cambium boundary and on the characteristic length, *λ.* Since *λ* was assumed to be constant (because the diffusion coefficient and decay rate of signal D are themselves constant), the width of the cambial zone was entirely driven by the concentration of signal D at the cambium boundary.

Regarding the enlargement zone, the width of the gradient of signal G was the main driver. For a given value of decay rate, this width increased with the height of the concentration peak, which in turn depended on the concentration of signal G at the cambium boundary and on the number of polar transporters in dividing cells. Since the number of transporters was assumed to be proportional to the local concentration of signal G, the width of the enlargement zone was entirely driven by the concentration of signal G imposed at the cambium boundary.

The patterns of variations that we imposed on the concentrations of signals D and G at the cambium boundary, as described above, lead to the variations in cell numbers in the cambial and enlargement zones represented in Fig. 4a. Comparison with experimental data from Cuny et al. (2014) displays good agreement across the growing season (Fig. 4b). This supports that developmental zonation can be adequately controlled by the cross-talk between two biochemical signals.

**Figure 4:**
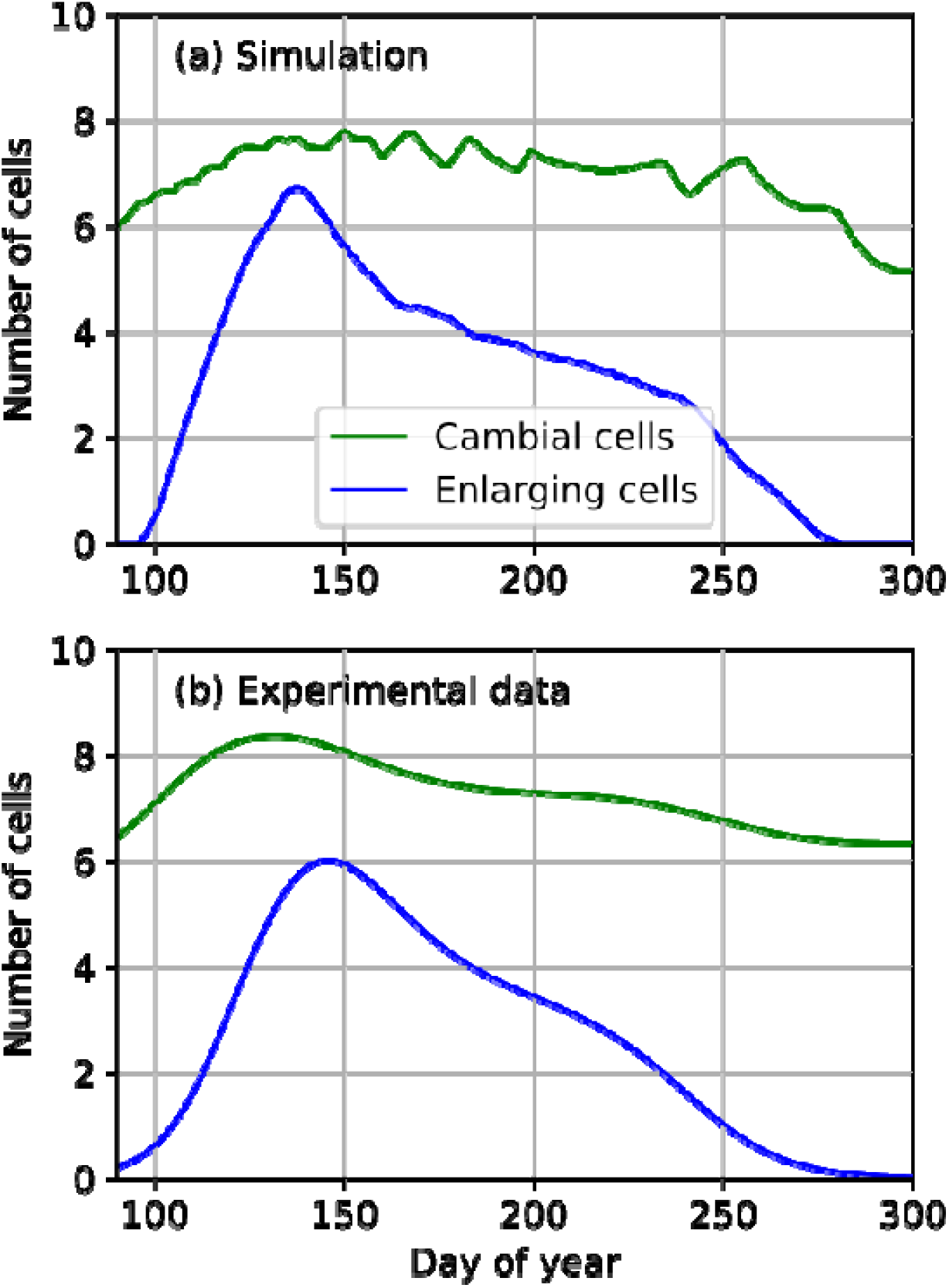
Evolution of the number of cambial and enlarging cells over a growing season. (a) As simulated by the XyDyS model. (b) From observations on silver firs (*Abies alba*) in the Vosges Mountains (France) reported in Cuny et al. (2014).

### The cross-talk between signal D and G engenders a realistic pattern of stem radial growth

The total growth rate of the cell file was directly related to the total quantity of signal G in the tissue. Three factors determined this quantity: 1) The concentration of signal G imposed at the cambium boundary; 2) the number of cells contributing to the polar transport of signal G (i.e. the number of dividing cells), controlled by the gradient of signal D; and 3) the amount of polar transporters in each of these cells (*p_i,i+1_*), which was itself directly proportional to the local concentration of signal G. As a consequence, the global growth rate of the cell file was controlled by the concentration of signal G at the cambium boundary and, to a lesser extent, by the boundary concentration of signal D.

With our hypotheses for the changes in the concentrations of signals D and G at the cambium boundary, the simulation resulted in the cumulative growth curve shown in Fig. 5a. It can be compared with measurements made on Scots pine *(Pinus sylvestris)* by Michelot et al. (2012), and displayed in Fig. 5b. In our simulations, we did not try to match the final cumulative growth, which depends on many factors, so only general shape of the curves should be compared. Although the agreement is not perfect, the simulated curve reproduces qualitatively the slow start, the progressive acceleration, the stable linear part, and the final progressive cessation of growth.

**Figure 5:**
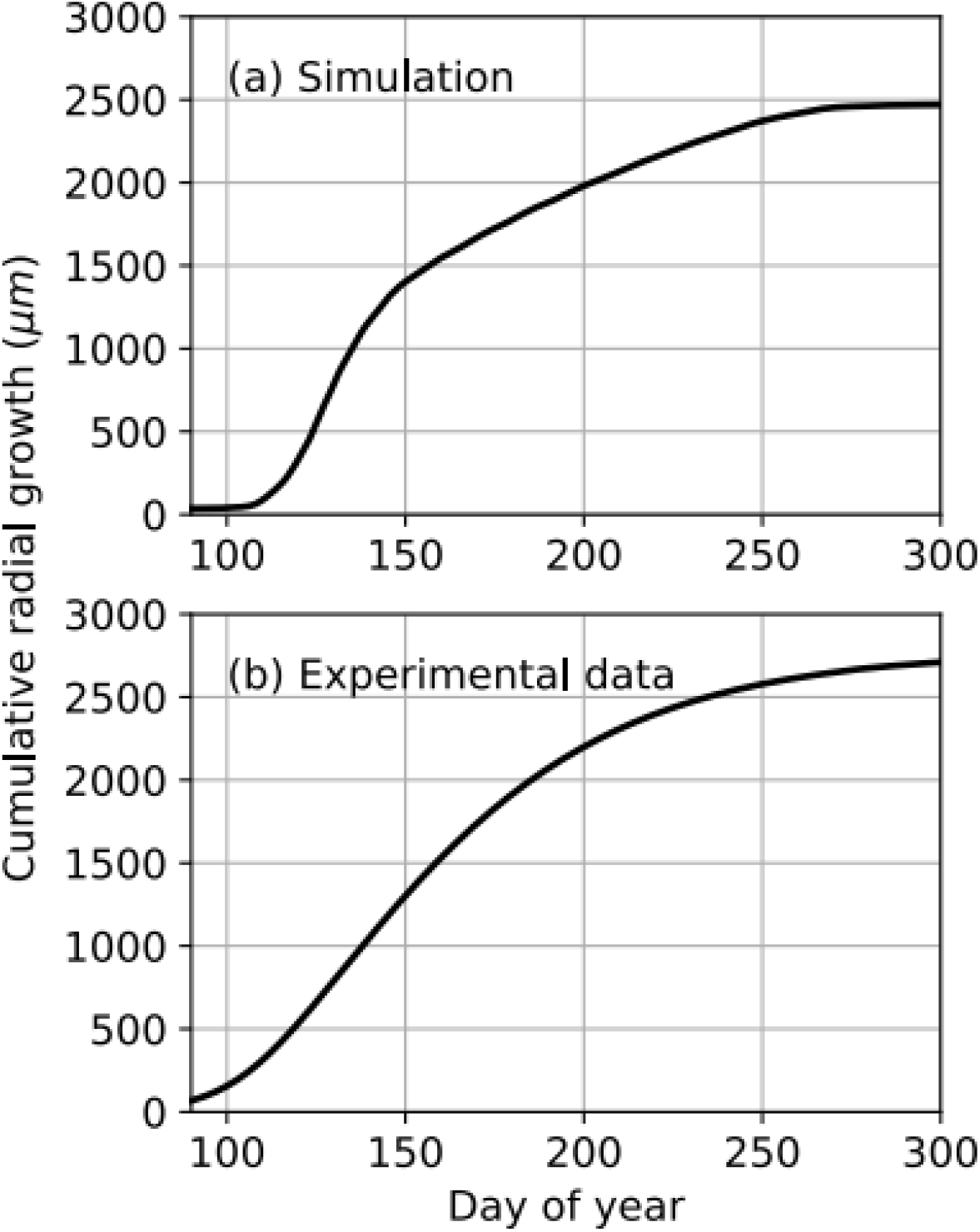
Cumulative radial growth of a tree ring. (a) As simulated by XyDyS; and (b) as fitted from microcore measurements on Scots pines *(Pinus sylvestris)* growing in Fontainebleau forest, close to Paris (France) and reported in Michelot et al. (2012).

### The cross-talk between signal D and G engenders a realistic tree-ring structure

We found that the final size of each tracheid was proportional to the height of the concentration peak of signal G at the time the cell lost its ability to divide. Indeed, the higher the peak is when the cell moves to the enlargement phase, the more signal G is available to the cell for this phase. The height of the peak depends in turn on the concentration of signal G on the cambium boundary and on the magnitude of active polar transport. The strong supply of signal G at the beginning of the growing season resulted in large earlywood cells. The progressive decrease in auxin supply during the progression of the growing season leaded to transition wood and, finally, narrow latewood cells (Fig. 6a).

**Figure 6:**
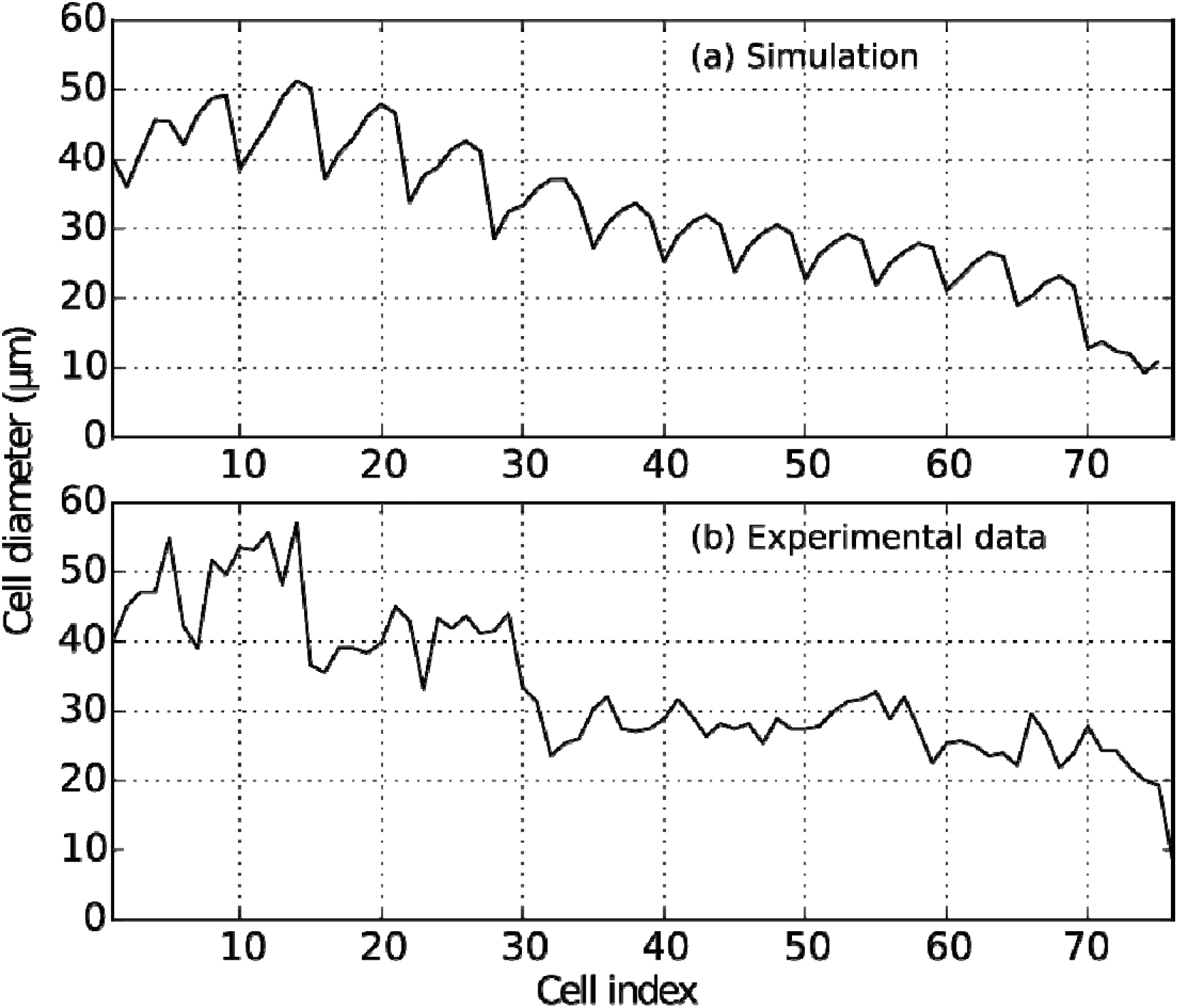
Evolution of tracheid radial diameters along a mature tree ring. (a) As simulated by XyDyS; and (b) as measured on a microcore of Scots pine *(Pinus sylvestris)* growing in the Vosges Mountains (France). Data courtesy from Henri Cuny.

The previous implementation of the morphogenetic-gradient hypothesis in a dynamical model predicted unrealistic regular spatial oscillations of high amplitudes in final cell sizes (Hartmann et al. 2017). Here, the size-dependence between auxin concentration and growth rates introduced in Eq. 9 greatly alleviated this problem. This hypothesis did not produce smooth variations in cell sizes along a tracheidogram, but rather moderate-amplitude irregularities that can also be found in experimental data (Fig. 6b).

## Discussion

In a previous work (Hartmann et al. 2017), we have shown that the morphogenetic-gradient hypothesis was not compatible with the anatomical structure of conifer tree rings. In the present article, we proposed a new model involving two biochemical signals. The first signal is associated with cell division and could be identified as the peptide TDIF, which is known to enter the cambium from the phloem and to be involved in vascular stem cell maintenance (Hirakawa et al. 2008; Etchells et al. 2015). Another candidate for this first signal is the plant hormone cytokinin, whose regulatory effect on cambial activity has been reported in aspen (Nieminen et al. 2008). The second signal is associated with cell growth and identified as auxin. Indeed, auxin is known to stimulate cell growth in many tissues, including stems (Perrot-Rechenmann 2010), and to inhibit secondary cell wall deposition (Johnsson et al. 2018).

Our model reproduced the main features of intra-annual dynamics of conifer wood formation over a growing season, i.e. the shape of the radial growth curve, the temporal evolution of differentiation zones, and the final anatomical structure of the tree ring in terms of tracheid radial diameters. It also provided an explanation for the pattern of auxin distribution in the developing xylem. The final radial size of cells was controlled by the supply of auxin to the cambium. Such a control was not possible with the classical morphogenetic-gradient hypothesis. It became possible by introducing two new hypotheses in XyDyS2: a decoupling of cell growth from division through the introduction of a second signal, plus a feedback of auxin on its own transport. With these new hypotheses, the final radial diameter of a tracheid was essentially set by its auxin content at the time it exits the cambial zone. This result supports the idea of hierarchical control proposed by Vaganov et al. (2011).

Our assumption that auxin polar lateral transport plays a significant role in wood formation is based on an experimental study on aspen by Schrader et al. (2003). In particular, they observed higher expression of PIN genes in dividing xylem cells than in expanding ones. This is why we assumed that PIN proteins responsible for lateral auxin transport are only present in dividing cells. However, there is no spatially-resolved direct measurement of the concentration and localisation of PINs in the cambium. Our hypothesis that PINs are polarised towards the xylem hence remains speculative. Another crucial hypothesis of our model is the auxin-dependence of PIN synthesis. Such auxin-dependence of PIN synthesis is strongly supported by experiments on apical meristems (Vieten et al. 2005), but so far there is no direct evidence of it in the cambium. Further experimental works are thus needed to get a better understanding of polar auxin transport in the developing xylem and assess our hypotheses.

In our previous modelling work, we reported large oscillations of final cell sizes along a simulated tree-ring (Hartmann et al. 2017). We showed here that these oscillations can be strongly attenuated by assuming that the growth response of cells is size-dependent. This is based on the biological idea that larger cells have a lower density of DNA in their cytoplasm (provided there is no endoreplication), and thus have a lower capacity to sustain growth. This hypothesis is supported by the works of Mellerowicz and Riding (1992) who did not find any endoreplication in *Abies balsamea* cambium. Further support for weaker growth response in larger cells comes from observations in sepal epidermis, where smaller cell lineages grow faster than larger ones (Tsugawa et al. 2017). This results in a homogenization of cell sizes. Moreover, in the shoot apical meristem of *Arabidopsis thaliana*, Willis et al. (2016) found that, after an asymmetrical division, the smallest daughter cell grows at faster rate than the largest one.

Although attenuated, fluctuations in final cell sizes were still present in our simulations. They were, however, similar in amplitude to fluctuations observed in actual tracheidograms. It is interesting to note that these oscillations are completely determined by the mechanisms behind the growth dynamics of the developing xylem tissue, without any explicit stochastic component. Numerous cellular processes involve stochastic component (Meroz and Bastien 2014; Meyer et al. 2017), and this aspect should be also investigated in the future. Nevertheless, our results underline that not all heterogeneities in cell features are attributable to stochastic processes.

We used a purely deterministic criterion for division, based on a cell size threshold. This assumption is supported by the probable existence of a cell size checkpoint at the G1-S transition (Schiessl et al. 2012). Moreover, analyses of cell size distribution along the growth zone of developing roots (Beemster and Baskin 1998) and leaves (Fiorani et al. 2000) suggest that all the cells in a given meristem divide in half at the same length. However, the critical-size criterion is likely to be essentially a first-order approximation. In the shoot apical meristem, Willis et al. (2016) found that it could not fully account for the cell-cycle statistics observed. Future modelling works could explore whether introducing stochasticity here can better reproduce wood anatomical structure.

We focused on biochemical signals to model wood formation, with no explicit mention of environmental factors. In reality, the inputs of our model, i.e. the supplies of signals into the cambium, are related to developmental and environmental conditions. These relationships are not known exactly, and tree-scale models are needed to connect signal sources to sinks. Moreover, temperature and water status also alter the capacity of cells to respond to signals. Here we made the implicit hypothesis that environmental conditions were not limiting. Further developments of our model could consider how wood formation dynamics is affected by water stress, which can be a limiting factor at least in the xeric area (Cabon et al. 2020a). Ignoring temperature effects also limits the scope of our model. For instance, we do not model the timing of the onset of cambial activity, which is likely to be triggered by temperature (Begum et al. 2012; Delpierre et al. 2019; Cabon et al. 2020b). Similarly, growth cessation in autumn may involve responses to day length (Baba et al. 2011), temperature (Begum et al. 2016), or even drought (Ziaco et al. 2016, Cabon et al. 2020b). Finally, wind-induced mechanical strains have been proved a major driver of wood growth rate (Bonnesoeur et al. 2016) and of wood anatomy (Roignant et al. 2018).

The final phases of xylem cell differentiation, i.e. secondary wall formation and programmed cell death, involve many biochemical processes. However, they may not be controlled by an additional signal. It has been observed that the amount of secondary wall material is about the same in each mature xylem cell along a tree ring, except for the very last latewood cells (Cuny et al. 2014). This observation could be used to deduce secondary wall thickness from cell size. Besides, temperature seems to play little role in wall thickness, since forming tracheids compensate a decreased rate of differentiation by an extended duration, except for the last cells of the latewood (Cuny et al. 2018).

Here we considered that the spatial organisation of the cambium relies only on biochemical signals. However, it is possible that the mechanical pressure exerted by the bark is also involved in cambial organisation, by setting a radial polarity field (Yeoman and Brown 1971). Mechanical signals are known to be essential in the dynamics of the shoot apical meristem, especially in the boundary region, where cells divide periclinally (Louveaux et al. 2016) just as in the cambium. While biochemical signals are likely to play a major role in controlling cell differentiation, division, and growth rate during the growing season, mechanics probably also provides cues to cambial cells. Future, more advanced models of cambial activity and wood formation should embrace both biochemical, environmental and mechanical signalling.

## Supplementary Data

S1 Video **Video of the simulation.**

## Acknowledgment

FPH thanks Mélanie Decourteix for helpful discussions.

